# Nervous system mediated cardiac arrest in mangrove killifish

**DOI:** 10.1101/2025.02.03.636252

**Authors:** Rosalie Herbert, Alexander David Corbett, Tetsuhiro Kudoh

**Author notes:** Contact addresses.

## Abstract

The mangrove killifish, *Kryptolebias marmoratus*, is a highly unique vertebrate species, known for its unusual sexual systems. It is one of two known self-fertilizing simultaneous hermaphroditic vertebrates, alongside the related *Kryptolebias ocellatus*^1,2^. While individuals can be born as primary males, most are born as hermaphrodites^3^ and some turn into secondary males later in life^4^. *K. marmoratus* is exceptionally tolerant of harsh and variable environmental conditions. This includes emersion as well as widely ranging salinities and temperatures^5–7^. Embryos can enter diapause, a state of arrested development, when conditions are unfavourable^8,9^.

---

Intentional heart stopping by animals is an exceptionally rare phenomenon. Stopping or significantly reducing the heart rate is also known to occur in some animals when in a state of dormancy such as hibernation or diapause^10,11^. However, an abrupt and complete heart-stopping behavior that occurs frequently, causes no harm, and is followed by full recovery, has not previously been reported.

We have found a unique ability in *K. marmoratus* embryos of heart stopping in response to subtle physical disturbances (HSPD), with fast and full recovery. Here, we investigate this behaviour for the first time. This ability begins in the later stages of embryonic development, prior to hatching/diapause. The behaviour of HSPD is then retained during diapause. Anaesthetic treatment with MS222 inhibited the HSPD ability, suggesting the behaviour is not autonomous to the heart but regulated by the nervous system. These results provide new insight into adaptive responses to environmental stressors and contribute to establishing this species as a model organism.

## Results

### Heart stopping and recovery

The heart of the *K. marmoratus* embryos started beating at 3 days post fertilization (dpf), as previously reported^12^. At the later stage of embryonic development from 9dpf, a peculiar heart stopping in response to physical disturbance (HSPD) was observed. The physical disturbances that triggered HSPD included gentle touching of the chorion (Supp. video 1) or by tapping the petri dish containing the embryos (Fig. 1). At 9dpf, stops were short, with full recovery averaging 7.78 seconds. From 9dpf to 14dpf, the duration of heart stops and the recovery time increased (Fig. 2).

**Figure 1.**
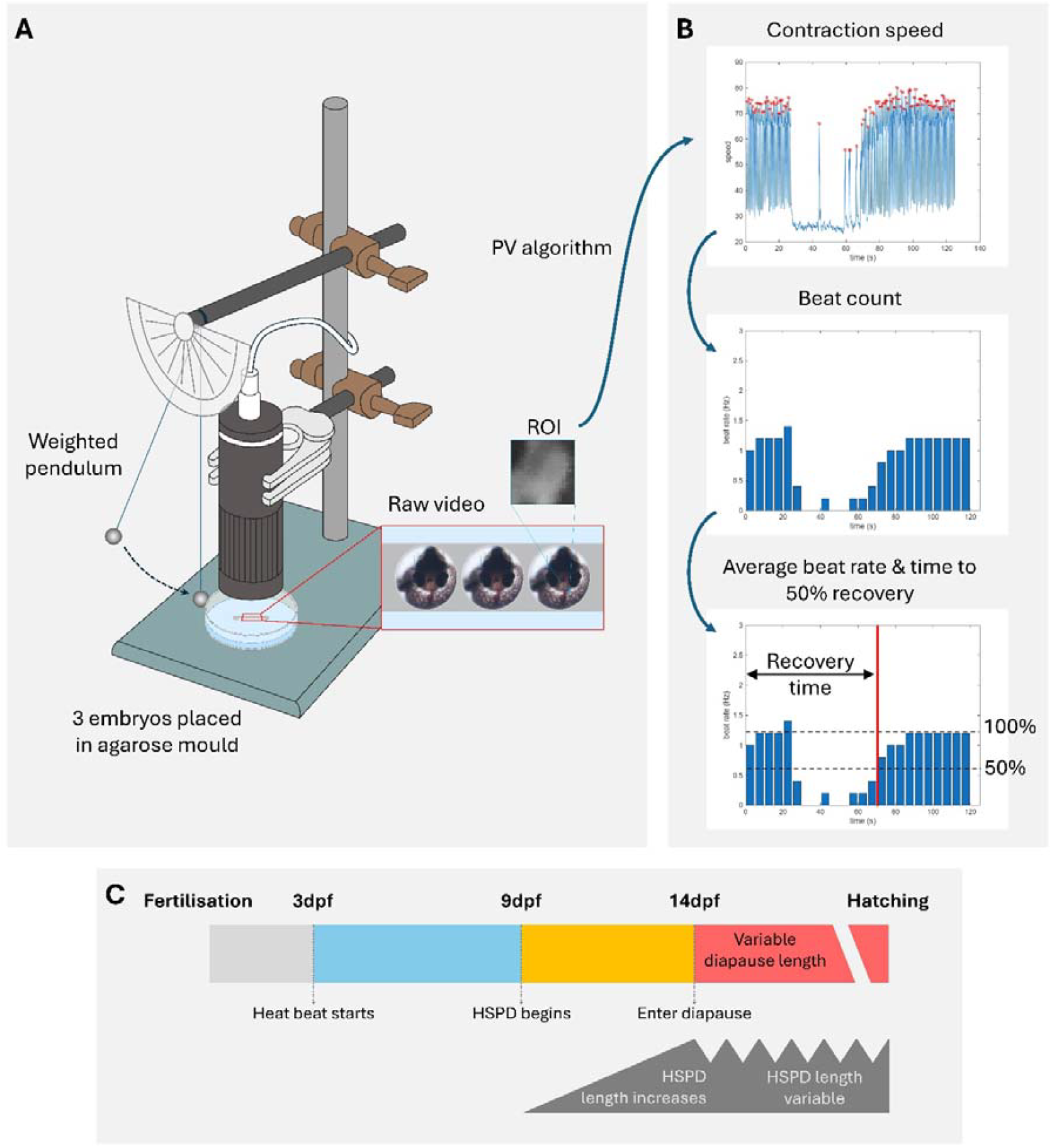
Summary of methods and results. (A) Set-up: embryos were placed in an agarose mold under a microscope, and HSPD was induced and filmed using a pendulum-based mechanical stimulation setup. (B) Data analysis: from the recordings, a region of interest (ROI) was selected over a single heart chamber. The Pixel Variance algorithm was performed to find contraction speed, and beat count was calculated (example shown for 12 dpf). From this, the baseline pre-stimulus average heart rate, and time to 50% recovery after HSPD were measured. (C) Summary of results: key stages in the development of HSPD from fertilization to hatching (top); grey panel (bottom) illustrates that the duration of HSPD increases from 9 to 14dpf and becomes variable during diapause.

**Figure 2.**
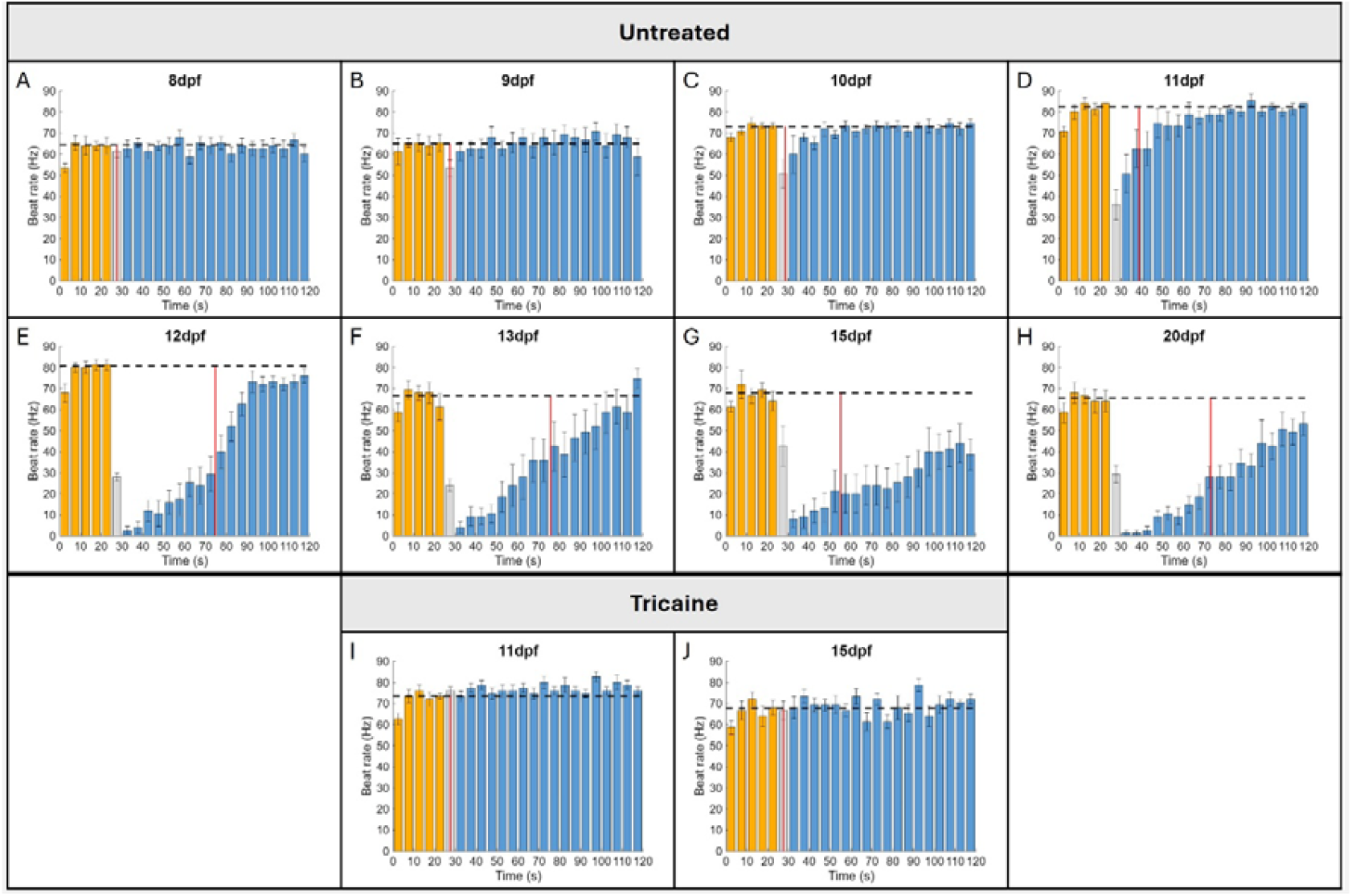
HSPD starts from 9dpf and increases in duration throughout development, but is inhibited by tricaine treatment. Heart rate was measured at 8-13, 15 and 20dpf for untreated embryos (A-H) and at 11 and 15dpf for tricaine-treated embryos (I-J) at 5-second intervals over 120s movies. At 25s, stimulation was applied, inducing HSPD. Untreated embryos stopped their heart beat in response to the stimulation, but tricaine-treated embryos lost this ability. Orange bars show heart rate before the stimulus at 25s, from which the baseline pre-stimulus average heart rate was calculated, indicated by the black dotted line. Blue bars show post-stimulus heart rate. Red lines mark the average time at which heart rate reached 50% of the pre-stimulus rate. The time points indicated by grey bars (27.5s) were not included in recovery time due to noise caused by embryo movement following the stimulus. Error bars show SEM.

Around 13dpf, embryonic development is complete, and embryos can either hatch or enter diapause^12^. Most embryos in our experiments entered diapause, delaying hatching. The HSPD ability was retained during diapause. However, once embryos entered diapause (∼14dpf), lengths of heart stops and recovery periods became more variable (Supp. fig.1). While embryos were responsive to physical stimuli, light stimulation (light on/off) did not induce HSPD (data not shown). During HSPD, the heartbeat ceased, but other movements, including those of the eyes, mouth and gills did not stop. This indicated that HSPD does not cause complete paralysis of the body (Supp. video 2). Embryos were highly sensitive, as gently touching the chorion with the tip of a tweezer induced HSPD and stops could exceed 60 seconds (exemplified in supp. video 1).

To determine whether HSPD was autonomous to the heart tissue or regulated by the nervous system, the effect of tricaine was tested. Tricaine blocks sodium ion channels in nerve cells, preventing initiation or transmission of signals. Treatment was applied to both developing (11dpf) and diapausing (15dpf) embryos. In all cases, tricaine-treated embryos lost HSPD abilities (Fig. 2I-J). This suggests that HSPD is not autonomous in the heart but is mediated by a signal from the nervous system. With tricaine treatment, pre-stimulus heart rates were reduced in 11dpf embryos (mean difference 8.66bpm, p=0.0103), but heart rates of 15dpf embryos were not significantly affected (p>0.05).

To examine if the response of HSPD depends on the level of physical stimulation, quantitatively reproducible levels of stimulation were given to embryos using a pendulum-based mechanical stimulation setup (Fig. 1). The data showed that for all embryonic stages tested (10 to 14dpf), the length of heart stops was stimulus-dependent, with stronger stimuli inducing longer heart stops (Fig. 3).

**Figure 3.**
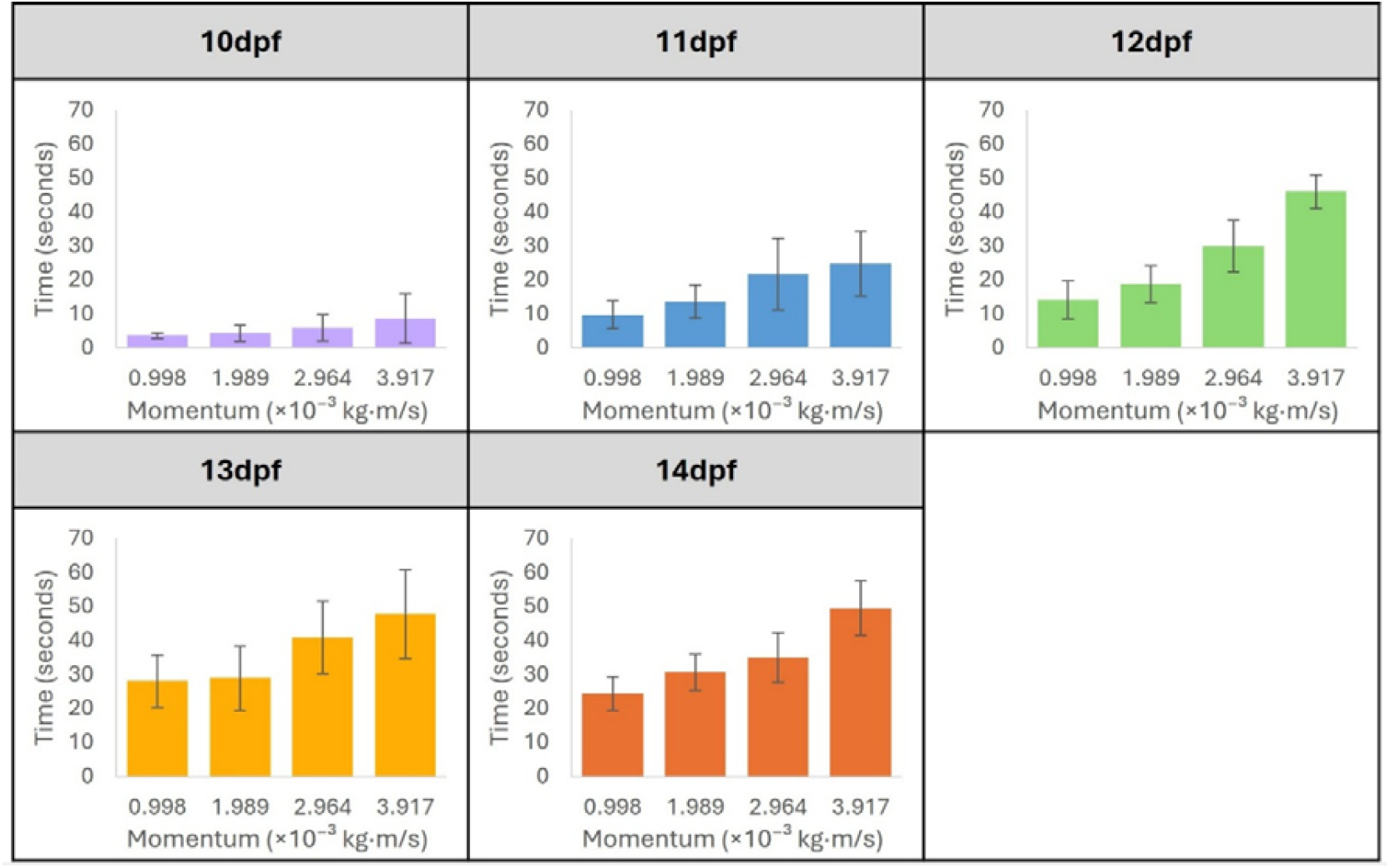
Effect of Stimulation Intensity on duration of HSPD. A range of physical stimuli with varying momentum were applied to *K. marmoratus* 10-14dpf embryos (n=9 per age group). With increasing stimulation momentum, the length of heart stops increased, measured by time from the stimulus to the first heartbeat after HSPD. Error bars show standard deviation.

Baseline pre-stimulus heart rates increased throughout development, peaking at 11-12dpf, before declining at 13dpf. Heart rates remained reduced throughout the diapause period, albeit with high variability (Supp. fig. 3). Diapause heart rates were significantly lower than at both 11 and 12dpf (mean differences 12.38 and 10.71bpm, p=0.0022 and 0.0082, z=3.0512 and 2.6427, respectively).

### Heart stopping and restarting occurs in the sinus venosus

In *K. marmoratus* embryos, individual contractions of the sinus venosus, atrium and ventricle were clearly distinguishable and could be measured independently (Fig. 4). Stimulation caused the heart to stop immediately, the last beat always ending with the ventricle contracting (Fig. 4E). During HSPD, the sinus venosus and atrium remained filled with blood, as in diastole, and passive flow into the ventricle was often observed. The first beat after HSPD consistently began with contraction of the sinus venosus (Fig. 4F). These results indicate that the order of chamber contractions is always sinus venosus, atrium, ventricle.

**Figure 4.**
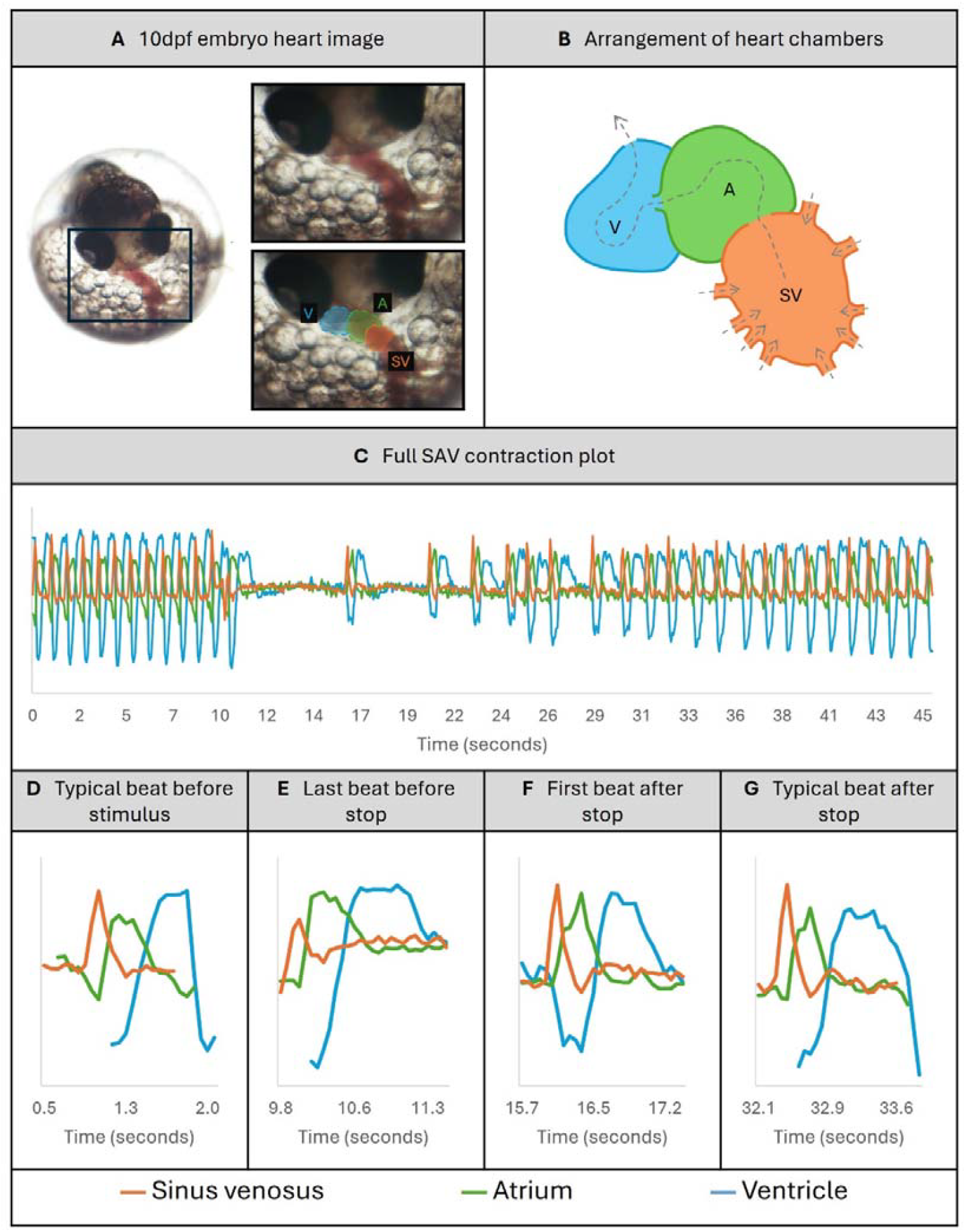
Heart chambers and contraction plots. (A) Image of *K. marmoratus* embryo at 10dpf, once HSPD ability is established. SV (sinus venosus) is indicated in orange, A (atrium) in green, V (ventricle) in blue. (B) Arrangement of chambers, arrows indicating the direction of blood flow. (C) A full SAV (Sinus venosus, Atrium and Ventricle) contraction plot from a 45-second movie of a 12dpf embryo. Physical stimulation was applied at ∼10s, inducing HSPD. Below, individual beats are highlighted: (D) a typical beat from before the stimulus, (E) the last beat before HSPD, (F) the first beat after HSPD and (G) a typical beat after the heart rate recovers. Peaks indicate contractions for each chamber. Lines are coloured according to the chamber.

The time between each chamber contracting remained regular even during the recovery period after HSPD (Fig. 4). There was no significant difference in the timing between contractions of individual chambers before and after HSPD (p>0.25). Before HSPD, the average time after contraction of the sinus venosus for atrial and ventricular contractions were 0.23s and 0.62s, respectively. After HSPD, these were 0.25s and 0.65s. Together, these suggest the heart stop is always initiated by the sinus venosus.

## Discussion

### HSPD as a part of embryonic developmental process occurring before diapause/hatching

We have discovered a peculiar heart stopping behavior of mangrove killifish embryos in response to physical disturbance. This HSPD may occur frequently in the wild, considering cause can be attributed to even very slight physical disturbance. To our knowledge, this has not been reported in any other species.

Embryos consistently acquired the HSPD ability around 9dpf, with all embryos able to stop their hearts by 10dpf. The duration of stops increased until the end of development at 13-14dpf. This indicates that the acquisition and development of the HSPD ability is part of the highly synchronized embryonic development in *K. marmoratus*, and it occurs whether they enter diapause or not.

When embryos reach 13dpf, some embryos start hatching^12^. Therefore, from 14dpf onward, embryos that did not hatch likely entered a diapause - a state of arrested development where metabolic rate is significantly reduced, used to cope with stressful conditions^9^. Supporting this, baseline heart rates reduced at 13dpf and remained low during diapause (Supp. fig. 3). The initial reduction at 13dpf may be part of the preparation for entering diapause. As metabolism is reduced during diapause, the animal must prioritize which processes or functions energy is put into. This indicates that HSPD is of high importance for survival or coping with stressors during the diapause period.

Although metabolic rate is generally low during diapause, there can be wide variation between individuals^13^. This was evident in the reduced, but highly variable baseline heart rates in diapause embryos (Supp. fig. 3). Within this time there may also be pre-diapause preparatory phases or post-diapause recovery phases, during which metabolic rates can vary^9^. Additionally, the length of diapause is highly variable, previous studies reporting lengths ranging from 23 to 108 days^14^. These factors may contribute to the observed variation in HSPD recovery time in diapause-stage embryos compared to the clear pattern seen in 9-14dpf embryos, especially if recovery is linked to metabolic rate. This suggests that the development of HSPD is complete by 14dpf, and the variable “depth” of diapause may lead to variability in the level of HSPD.

Other killifish species (*Nothobrancheus furzeri* and *Austrofundulus limnaeus*) have been reported to reduce their heart rates that occurs as metabolism is reduced during diapause, sometimes to the extent they appear to stop completely^15,16^. However, in these species the heart stops without physical stimulation, and it occurs earlier on in embryonic development. In contrast, HSPD in *K. marmoratus* is an immediate, short-lived reaction to physical stimulation. It may be possible that the mechanisms to reduce/stop the heartbeat during diapause seen in *N. furzeri* and *A. limnaeus* may be exploited in the mangrove killifish to develop the ability of HSPD.

HSPD has not been reported in other fish species, suggesting that it may be an evolutionary survival strategy unique to *K. marmoratus*. It is conceivable that HSPD may have two advantages for embryos. Firstly, as mentioned above, HSPD is possibly a function associated with diapause. By being able to control heart rate, the embryo may be able to preserve energy in diapause. The second potential advantage of HSPD may be related to predator avoidance. *K. marmoratus* often inhabits crab burrows. During dry seasons they may remain in these small spaces with limited water supplies for weeks or months within a state of diapause. This could increase risk of predation by creatures such as small crabs. HSPD may reduce vibrations or electric pulses that could be detected by hair sensors of small crabs searching for food in these burrows. This was indicated by imitating the movement of a crab’s claw using the tip of a pair of tweezers and observing how this very small physical disturbance induced HSPD (Supp. video 1). Stopping the heartbeat to reduce metabolism and avoid predation may continue to be advantageous for juveniles and adults. Therefore, it would be interesting to investigate HSPD in hatched fish for future research. Alternatively, HSPD may not confer any adaptive advantages. It may be a result of linkage to other genes under selection, or the high levels of inbreeding and homozygosity in the mangrove killifish, which have arisen from selfing and may have led to the fixation of the trait^17^.

### Sinus venosus starts HSPD and restart of the heartbeat after HSPD

We observed that contraction of the sinus venosus was always followed by contraction of the atrium then the ventricle. When HSPD occurred, the ventricle was always the last chamber to contract and the sinus venosus was the first to contract. After HSPD, timings of these contractions of chambers within individual heartbeats remained the same as before HSPD, despite heart rate starting slower before recovering to the original rate. In mammalian hearts, the atrioventricular (AV) node delays the electrical signal from the atrium to ventricle – fish have an analogous structure within the AV junction to fulfil this role ^17^. There is also a delay between the sinus venosus and atrium contracting. In reptiles, the sinoatrial junction causes this delay^18^. Our results indicate that the key domain in which stopping and restarting occurs is the sinus venosus, and the other two chambers cannot stop or restart contraction on their own.

### HSPD is mediated by the neuronal circuit

Our results suggest that the sinus venosus is the key domain of the heart determining the initiation of HSPD and release from it. In fish, pacemaker cells are located in a ring-like shape at the sinus node, initiating contraction^19^. Therefore, it seems likely that the pacemaker cells are involved in regulation of HSPD. Vertebrate hearts are myogenic, generating their own rhythm independent of the brain. Pacemaker cells within the heart itself initiate the electrical impulse. However, results from tricaine experiments suggest that the heart stopping response is also dependent on neuronal function. Tricaine blocks sodium ion channels in nerve cells, preventing them from initiating or transmitting signals. Treating embryos with tricaine completely prevented HSPD. Heart rate can be modulated by the autonomic nervous system (ANS); sympathetic neurons enhance heart rate while parasympathetic neurons reduce it. HSPD may be mediated by activation of parasympathetic neurons. However, it is uncertain whether this could apply to embryos as it depends on whether the ANS is sufficiently developed at this stage. For example, medaka embryos, despite development of autonomic nervous fibers as embryos, may not be able to regulate heart rate until one month post-hatching^20^. Therefore, further research is needed to elucidate the molecular and cellular mechanisms of HSPD.

## Ethical declaration

All experiments and methods were performed in accordance with the relevant guidelines and regulations of UK Home Office with the license number PP4402521. All methods are reported in accordance with ARRIVE guidelines. All experiments and methods approved by the Animal Welfare and Ethical Review Board (AWERB) at the University of Exeter.

## Resource availability

### Materials Availability

No new reagents were generated in this study.

### Data and Code Availability

The research data and Pixel Variance code supporting this publication are openly available from the University of Exeter’s institutional repository (ORE) at https://doi.org/10.24378/exe.5506. A description of the Pixel Variance algorithm is given in reference ^21^.

## Acknowledgements

TK is funded by NC3Rs project grant (NC/X001121/1). RH was funded by University of Exeter – Employer Engagement & Student Employment (EESE). We thank Sulayman Mourabit, Thomas Cosgrove, Rachel Osman, Clare Bushell, Ed Grinsted, Maryia Fralova and Matt Winter for their support. We also thank Aquatic Resource Centre staff for fish husbandry and to Biosciences teaching lab staff for sharing the lab space and equipment.

## Author Contributions

Conceptualization TK, Data curation RH, Investigation RH, Methodology TK, ADC, RH, Writing – original draft RH, TK, Writing – review & editing RH, TK, ADC

## Declaration of interests

The authors declare no competing interests.

## STAR Methods

### EXPERIMENTAL MODEL AND STUDY PARTICIPANT DETAILS

#### Mangrove killifish source and husbandry

The mangrove killifish *Kryptolebias marmoratus* wild type Hon9 line^22^ was sourced and maintained in the Aquatic Resource Centre at the University of Exeter. These fish were kept in a recirculation system under artificial photo periods of 14 hours of light to 10 hours of darkness. All fish were hermaphrodites.

#### Embryo collection/incubation

For this study, the *K. marmoratus* Hon9 line^22^ was used. Embryos were incubated at 28°C in 14.6ppt artificial brackish water. Water changes were made every other day. Embryos were aged according to the Mourabit *et al*., 2011 staging reference^12^. Embryos used for this study were ages 7 to 20dpf.

### METHOD DETAILS

#### Imaging & filming heart stopping

To administer controlled and reproducible physical stimuli to embryos held in a petri dish, a pendulum-based setup was assembled. Agar moulds to hold embryos were prepared using 1% molecular-grade agarose in deionized water in a 6cm petri dish, with a duplex cable wire to form a well to hold embryos. Three embryos were placed in the mould with 14.6ppt water. and positioned under a microscope at the base of a clamp stand (Fig. 1). A small steel ball (4.09g) was strung from a horizontal bar 20cm above the edge of the petri dish. Embryos were left to settle for 5 minutes to allow for recovery from HSPD that may have occurred from moving embryos before filming. The pendulum could be drawn back to a specified angle and released, hitting the petri dish and stimulating HSPD.

First, to assess whether the size of shock impacted HSPD, embryos aged 10-14dpf (*n*=9 for each age) were filmed under the described set-up and stimulated using a range of release angles (10-40°). Having tested this, a release from an angle of 20° was selected to use for the main analysis of HSPD, equating to a momentum of 1.989×10^− 3^ kg·m/s. This stimulus induced HSPD without causing embryos to shift within their chorions, which sometimes occurred at higher momentums, making image analysis difficult. For this main analysis, embryos were filmed for 120s, 25s into the movie the pendulum was released to hit the petri dish, inducing HSPD. Embryos aged 7-20dpf were used (*n*=9 for each age).

To assess whether embryos were also responsive to light stimulation, three embryos were placed under the microscope with the light off. Embryos were filmed for 60s, and 25s into the movie the microscope light was switched on.

Movies were recorded using the digital microscope Jiusion 2K HD 2560x1440P USB Digital Microscope VirtualDub^23^ in .avi format. For anatomy imaging, Nikon SMZ1500 and Digital Sight DS-U2 camera were used.

#### Tricaine treatment

Nine embryos for ages 11dpf and 15dpf each were initially treated with 0.08% tricaine/14.6ppt artificial brackish water for 30 minutes before starting filming. In some cases, this dose had no noticeable effect (probably due to their hard/thick chorion), so the dose was increased in 0.04% increments until tricaine was having an observable effect (lack of jaw and fin movement). The final doses used were 0.08% for 11dpf, and 0.16% for 15dpf. Embryos were then filmed using the same methods used for the main analysis.

### QUANTIFICATION AND STATISTICAL ANALYSIS

#### Influence of shock strength on HSPD

Using movies stimulating embryos with varying pendulum release angles, the influence of increasing stimulus momenta on HSPD was quantified. The time from administration of the stimulus to the first heartbeat following HSPD was timed. Averages and standard deviation were calculated for each age and release angle.

#### Heart rate, stopping and recovery analysis

Videos for the main analysis of HSPD, as well as tricaine treatment experiments and response to light stimulation were analysed. ImageJ^24^ was used to select the ROI over a single chamber, convert to 8-bit greyscale and export to a TIF image sequence. This image sequence was analysed using a ‘Pixel Variance’ algorithm which has been previously applied to the contraction of cardiomyocyte monolayers^21^. In short, the algorithm calculates the variance in the pixel value across frames captured over a known time interval. This time interval corresponds to ∼40% of the beat period, i.e. 0.4 seconds for the resting 1 Hz heart rate, equivalent to two frames when sampling at 5.7 fps. Once the variance has been calculated on a pixel-wise basis, the value is averaged across all the pixels in the region of interest (ROI). A trace of this value over time provides a measure of the speed with which the pixel values are changing. For a single beat, there will be two peaks in the speed profile, corresponding to contraction and relaxation. The baseline value of the contraction speed varies between ROIs and samples due to the presence of local features and differences in average intensity. Peaks in the ‘contraction speed’ traces were then identified using the peak prominence (typically >50% of the peak height) relative to the baseline rather than hardcoding the absolute peak height.

Once the timing of the individual peaks had been identified, the number of peaks within five second windows (starting from time zero) were calculated and divided by two to obtain the number of complete heart beats per five second interval. This ‘beat count’ data, averaged over a five second window, was attributed to the midpoint of the window (i.e. 2.5, 7.5, 12.5 etc. seconds).

Baseline heart rate was calculated for each embryo (untreated as well as tricaine treated) using pre-stimulus beat count data from the first 25 seconds of the movies. Averages and standard deviation were then calculated for each age. For each embryo, the time taken for heart rate to recover to 50% of the pre-stimulus rate was measured. The first point of recovery considered was the 27.5 timepoint as the 25s timepoint was not an accurate representation, due to the noise in data following the mechanical stimulation at 25s which often caused the embryos to slightly rotate inside the chorion. For this reason, very short heart stops were not captured. Averages and standard deviation for time to reach 50% recovery were then taken for each dpf. The Mann-Whitney U-test was used to compare heart rates for each age in the “developing” category (7–13dpf) to the “diapause” category (14–20dpf). The Mann-Whitney U-test was also used to compare heart rates between untreated and tricaine treated embryos for individual ages.

#### Contraction profiles of individual chambers

ImageJ was used to select a ROI over the sinus venosus, atrium and ventricle separately and plot separate z-axis profiles. Data points were extracted and plotted on a single graph for each movie. Drift was removed using a Butterworth high-pass filter, with zero-phase delay.

For three embryos, the durations between contractions of individual chambers were measured. For each embryo, four measurements were taken before HSPD and four after. The Wilcoxon signed-rank test was then used for each embryo to compare the timing between chamber contractions before and after HSPD.

## Notes

https://doi.org/10.24378/exe.5506

### Competing Interest Statement

The authors have declared no competing interest.

### Summary of Updates

Improved stats. More quantitative method applied to give a physical shock.

